# Plant-environment microscopy tracks interactions of Bacillus subtilis with plant roots across the entire rhizosphere

**DOI:** 10.1101/2021.02.13.430456

**Authors:** Yangminghao Liu, Daniel Patko, Ilonka Engelhardt, Timothy S George, Nicola Stanley-Wall, Vincent Ladmiral, Bruno Ameduri, Tim J Daniell, Nicola Holden, Michael P MacDonald, Lionel X Dupuy

**Affiliations:** School of Science and Engineering, University of Dundee, Dundee, UK; Ecological Sciences, The James Hutton Institute, Dundee, UK; Department of Conservation of Natural Resources, Neiker, Bilbao, Spain; School of Life Sciences, University of Dundee, Dundee DD1 5EH, United Kingdom; Ingénierie et Architectures Macromoléculaires, Institut Charles Gerhardt, CNRS, University of Montpellier, ENSCM, Montpellier, France; Department of Animal and Plant Sciences, The University of Sheffield, Sheffield, UK; Northern Faculty, Scotland’s Rural College, Aberdeen, United Kingdom; IKERBASQUE, Basque Foundation for Science, Bilbao, Spain

## Abstract

Our understanding of plant-microbe interactions in soil is limited by the difficulty of observing processes at the microscopic scale throughout plants’ large volume of influence. Here, we present the development of 3D live microscopy for resolving plant-microbe interactions across the environment of an entire seedling growing in a transparent soil in tailor-made mesocosms, maintaining physical conditions for the culture of both plants and microorganisms. A tailor made dual-illumination light-sheet system acquired scattering signals from the plant whilst fluorescence signals were captured from transparent soil particles and labelled microorganisms, allowing the generation of quantitative data on samples approximately 3600 mm^3^ in size with as good as 5 μm resolution at a rate of up to one scan every 30 minutes. The system tracked the movement of *Bacillus subtilis* populations in the rhizosphere of lettuce plants in real time, revealing previously unseen patterns of activity. Motile bacteria favoured small pore spaces over the surface of soil particles, colonising the root in a pulsatile manner. Migrations appeared to be directed towards the root cap, the point “first contact”, before subsequent colonisation of mature epidermis cells. Our findings show that microscopes dedicated to live environmental studies present an invaluable tool to understand plant-microbe interactions.

## Introduction

The ability of plants and microorganisms to cooperate to capture soil resources underpins life in terrestrial ecosystems. In modern crop production systems, where these natural plant-microbe interactions have largely been replaced by artificial fertiliser input, it is thought that crop varieties may have lost the ability to maintain a diverse microbiome (Pérez-Jaramillo et al., 2016) and as a consequence the sustainability of the system has declined. Consequently, understanding of plant microbe interactions has become a major focus of research. Technological development has greatly expanded the knowledge of the microbial composition of soil: metabolomics details the chemical composition of organic material deposited by the root and high-throughput sequencing now describes the huge complexity of microbial communities associated with them (Widder et al., 2016). Soil habitats, however, are incredibly dynamic and structurally complex. The behaviour of the microbes inhabiting the inner structures of soil are equally complex, and to date current approaches have failed to provide mechanistic understanding of soil microbial dynamics (Zhao et al., 2014).

Since the discovery of cells and microorganisms, microscopy has constantly improved and modern microscopes are now able to solve problems of considerable complexity (Haseloff, 2003; Lane, 2015). However, live microscopy of plants within the biotic and abiotic environment remains complex and rarely achieved. Processes within the opaque world hidden within the soil structure are particularly difficult to monitor. Current microscopy methods applicable to soil are either destructive (Watt et al., 2006; Tuovinen et al., 2004), operate with samples of extremely limited volume and area (Deng et al., 2015), or oversimplify the role of the physical and chemical structure of the soil material (Massalha et al., 2017). Maintaining a viable undisturbed biological system is also a challenging condition to meet in the laboratory because processes occur both below and above ground, with different controls required for light, temperature, water and mineral content (Ramin Shamshiri et al., 2018).

The aim of this study was to build an “environmental microscope”, which we define as a live-sample imaging platform dedicated to the observation of physical and biological interactions that are relevant to the understanding of processes at environmental or system level. The platform we propose exploits recent advances in transparent soils, mesofluidics, and light-sheet imaging. It acquires fluorescence and scattering signals across the entire spatial domain surrounding a plant root, simultaneously combining all necessary controls for light, temperature and water content within the mesocosm. This study reveals previously unobserved phenomena of how bacteria colonise the rhizosphere, the region of soil surrounding plant root, and demonstrate the potential of the method to fill important knowledge gaps in environmental biology.

## Results

### Light sheet imaging for whole plant-environment microscopy

Observations were made from lettuce seedlings, a tractable plant for mesocosm studies, and *Bacillus subtilis* which is a well characterised rhizobacterium. Custom made chambers were assembled from glass slides and silicon parts to seal 4560 mm^3^ of transparent soil, water, nutrients, and atmosphere. The model system studied, therefore, was the entire environment supporting the lives of both plants and microbes (Figure 1 A).

**Figure 1.**
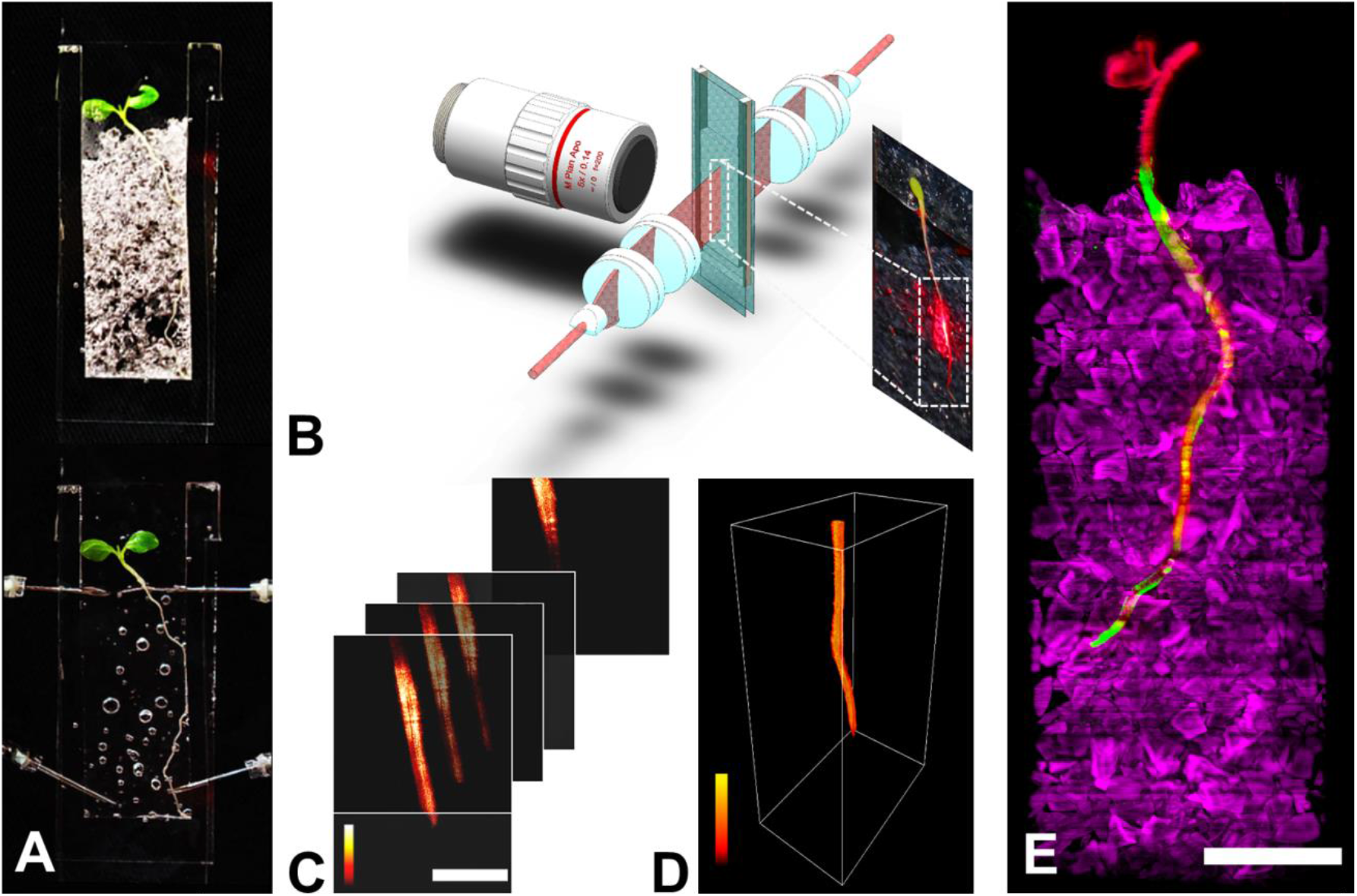
Live microscopy of the whole plant-environment. (A) Samples consisted of mesocosms filled with transparent soil and co-inoculated with lettuce plants and *Bacillus subtilis* (top). To perform imaging, the sample is saturated in refractive index matching liquid (bottom). (B) Light sheet microscopy (not drawn to scale) consists of a long focus homogenous light sheet generated using a Powell and two cylindrical lenses (scale bar 2 mm). The light sheet produces fluorescence and scattering signals captured by a long working distance objective. The mesocosm is immersed in refractive index matched solution, and is positioned and translated at a 45-degree angle within the light-sheet and detection arm. (C) Image data is acquired by translating the sample first in the horizontal plane, then vertically (scale bar 1 mm). (D) A complete volume dataset is stitched from a series of horizontal scans. (E) The microscope captured volume data of up to 3×60×20 mm^3^ and tracked the growth of entire seedlings (scattering, red), transparent soil particles (Rhodamine B fluorescence, magenta) and bacterial concentration (GFP-tagged micro-organisms *Bacillus subtilis*, green). Scale bar is 5 mm (E).

Acquiring biological signals from such volumes necessitates instruments that combine dedicated microscopy with adequate control hardware, and software. In this study, we showed Light Sheet Fluorescence Microscopy (LSFM) meets requirements for scale, throughput and integration with live mesocosm (Ovečka et al., 2018). The light sheet sectioned the sample optically with laser illumination optics, and the camera sensor captured fluorescence and scattering signals perpendicular to the plane of the light sheet (Figure 1 B). This was preferable to the use of condenser lenses and objectives with high numerical aperture (NA) for illumination and image capture because a high NA lens creates a shallow depth of focus (Dodt et al., 2007) and limited field of view, both of which are incompatible with imaging of large samples.

To achieve adequate throughput, and to limit the need for the stitching of multiple views, large field of view objectives (5× or 2× objectives with a field of view 2.4 mm or 6 mm respectively) were combined with a light sheet a centimetre in width and height. The light-sheet was generated with Powell lenses (Reidt et al., 2016) and series of cylindrical lenses that focused the sheet to the focal plane of the imaging objective with a thickness of 50 μm (from theoretical limit of 47 μm), a height of more than 5 mm, and a Rayleigh range (depth of focus) of at least 6 mm (Supplementary Information part 1&2).

Data from the entire mesocosm volume was successfully reconstructed (Figure 1 C&D) from custom made software that aligned the overlapping scans, corrected for artefacts from the imaging system, and produced a unique volume image containing signals from root, bacteria and soil particles (Supplementary Information part 3&4). The data generated by the microscope (Figure 1 E) was suitable for quantification of biological features by image analysis. The fluorescence from the bacteria was calibrated to predict cell density. Segmentation of the fluorescence signal from particles described how bacteria utilised the soil micro-structure while the scattering signal from the root measured the size and geometry of the root to accurately position bacterial population during growth, migration and colonisation on the root surface (Figure 2 and Supplementary Information part 5).

**Figure 2.**
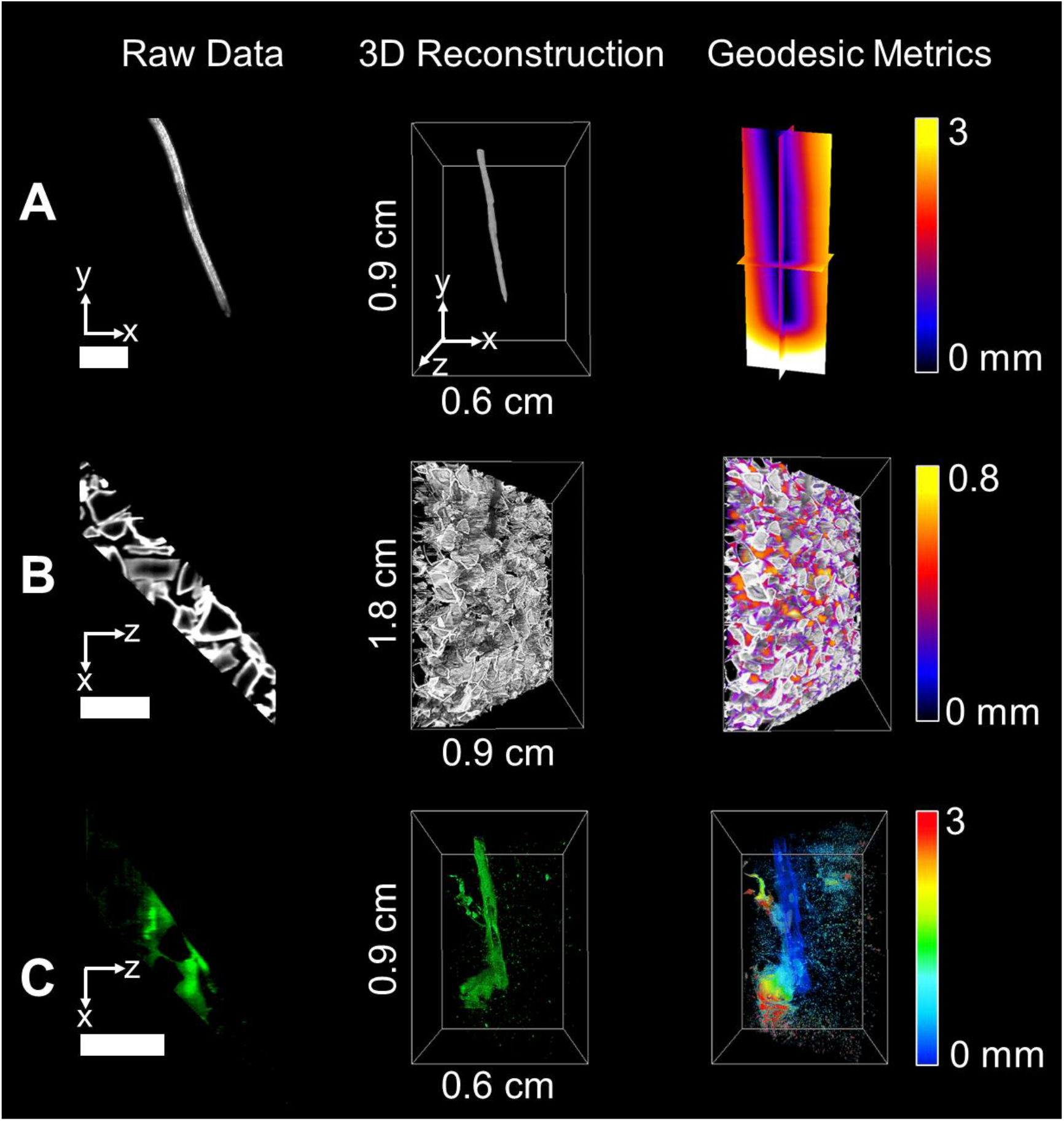
Quantification of root-soil-bacteria interactions. Image data from lettuce root (A), transparent soil particles (B), and GFP-labelled *Bacillus subtilis* (C). Processing of the data follows 3 steps. Raw data is acquired from the microscope (left). Cross sections are assembled into volume data through stitching and stacking (middle). Image processing is subsequently performed to quantify temporal and spatial patterns of biological activity in the pore space (right). The metrics obtained from the data include distance from the root surface, pore size and bacterial cell density. The scale bar represents a distance of 2 mm.

### Environmental microscopy resolves bacterial dynamics in the pore space

We observed movements of GFP marked *Bacillus subtilis* strain NCIB 3610 within the soil volume as well as interactions with the surface of growing lettuce roots. Quantification was performed every 30 minutes over 23 hours. The data acquired during those experiments was used to map the bacterial cell density in relation to the distance from the root surface, the distance along the root, and the size of soil pores. Significant bacterial colonisation was observed on six samples (out of 8). Two samples did not show bacterial colonisation and were not included in the study.

Overall, bacterial cell density estimated from pixel intensity was significantly greater when closer to the root surface. The phenomenon was particularly visible in the soil surrounding the base of the root (Figure 3 A, left), where the majority of bacterial cells concentrated within a radius of 1.5 mm from the root surface. By contrast, bacteria surrounding the root tip were observed within a radius of more than 3 mm from the root surface. Although the presence of bacteria was detected in all pore sizes, bacteria preferentially occupied smaller pores (<400 *μm*) of the soil (Figure 3 A, right).

**Figure 3.**
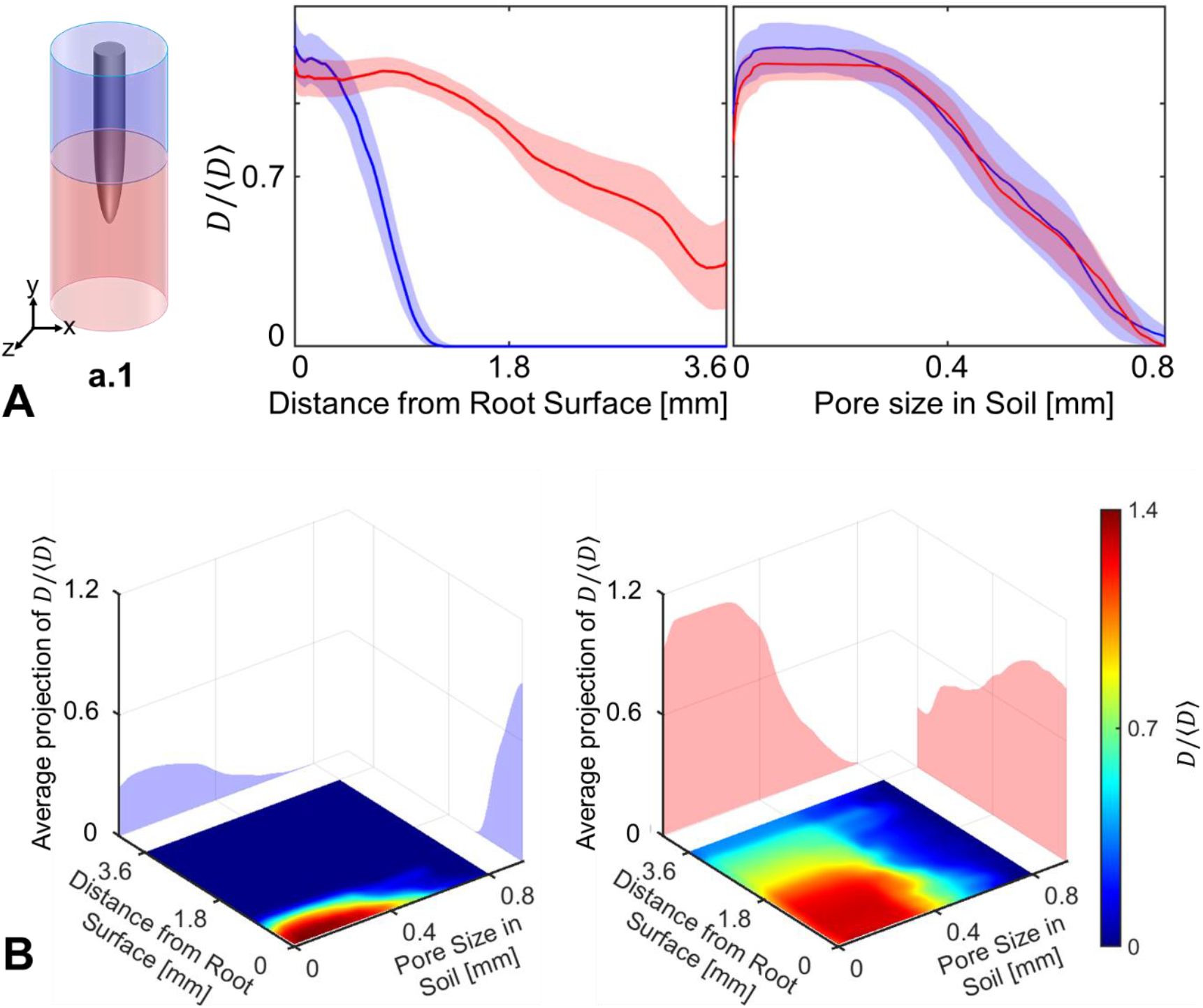
Utilisation of the pore space by *Bacillus subtilis* during colonisation. (A) The distribution of bacterial cell density varies as a function of the distance from the root surface. When bacteria are surrounding the basal region of the root (blue) they are present primarily in a radius of 1 mm around the root. When surrounding or in front of the root tip (red) bacteria are found within a radius around the root that is larger than 3 mm. There is little difference in the distribution of bacteria in the pore space. Data shown as mean ± SE, n=6. (B) The occupation of the pore space varies with the distance from the root. Bacteria tend to occupy larger pore spaces when further away from the root, however, the effect is more visible when bacteria are surrounding the basal region of the root (left) than near the root tip (right).

We observed a weak relationship between the pore size occupied by bacterial cells and the distance from the root surface. Plots of joint bacterial cell density distribution (Figure 3 B) showed bacteria closer to the root surface occupied the smaller pore spaced (<400 *μm*) in both the apical (left) and basal region (right) of the soil. On the contrary bacteria from the bulk soil occupied the pore space more evenly.

### Bacteria form hotspots and colonise the rhizosphere in pulses

Unlike growth in liquid culture, soil provides a physical support for bacterial attachment, but limits movement and confines cell activity to the pore microenvironment. To better understand how the pore space segregates the activity of *Bacillus subtilis* during root colonisation, a detailed analysis of the population dynamics was needed. Time-lapse data (over 23 hours) revealed increased bacterial cell density forming in specific regions of the soil, giving “hotspots” close to the root, or in the soil surrounding the root tip. Increased bacterial cell density was also observed on the root surface via attachment and / or biofilm formation (Figure 4 A). The location of bacterial hotspots varied significantly with time. Changes occurred more frequently near the root tip and hotspots appeared to stabilise on mature parts of the tissue, indicating attachment and formation of biofilms.

**Figure 4.**
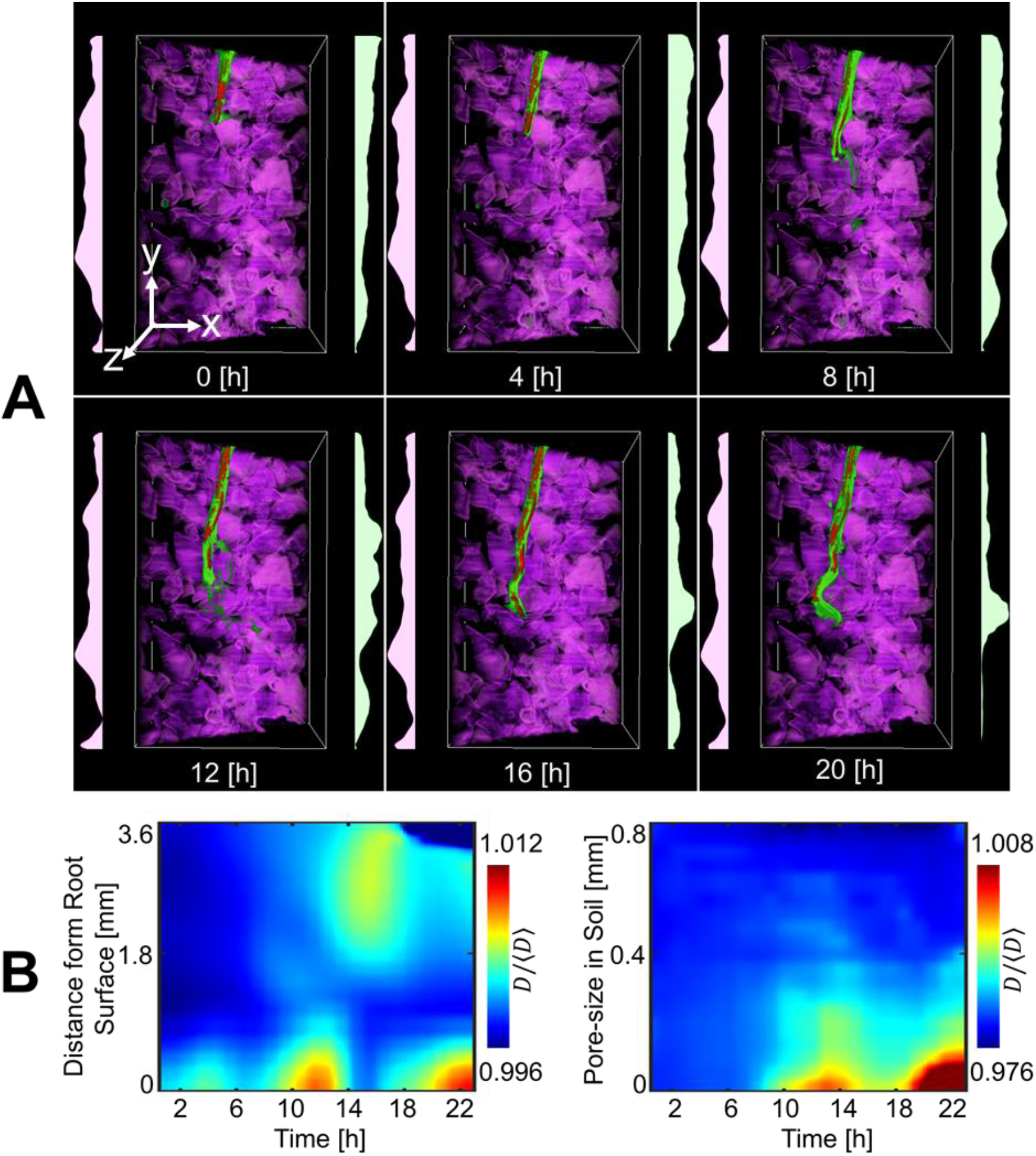
Time lapse data of the distribution of *Bacillus subtilis* in transparent soil surrounding lettuce roots observed for 20 hours after inoculation. (A) 3D visualisation of bacterial cell density reveals highly dynamic patterns due to interactions with the micro-structure of soil. (B) During colonisation, bacterial cell density increased in pulses, forming patches of bacteria close to the root surface (left). The increase in bacterial cell density occurs primarily in the smaller pore space (right).

Changes in bacterial cell density over time indicated that rhizosphere colonisation was pulsatile (Figure 4 B). Hotspots of bacteria appeared approximately five hours after inoculation in the smaller pore sizes (Figure 4 B, right), and was maintained for typically two to four hours following their appearance (Figure 4 B, left). Hotspots were observed at distances of more than 3 mm from the root surface, although the distance appeared to diminish during the course of the colonisation.

### Early interaction with the root cap may precede colonisation of the root surface

Large differences in bacterial cell density were observed between the overall increases in the bulk soil and the rhizosphere (Figure 5 A). The increase in mean bacterial cell density was first observed in the bulk soil, six hours after inoculation (Figure 5 A, red). Following a peak in bacterial cell density, the population of bacteria in the bulk soil subsequently reduced and reached a steady state. The increase in bacterial cell density in the rhizosphere was more gradual and reached a peak 10 to 14 hours after inoculation (Figure 5 A, blue). Fluctuations were observed following the peak of bacterial activity, but bacterial cell density remained high until termination of the experiment. The overall quantity of bacteria, calculated across the entire soil volume in the system, did not increase after reaching the peak concentration in the bulk soil (Figure 5 A, green). This indicates subsequent changes in bacterial cell density may largely be induced by bacterial movements through soil and along the root.

**Figure 5.**
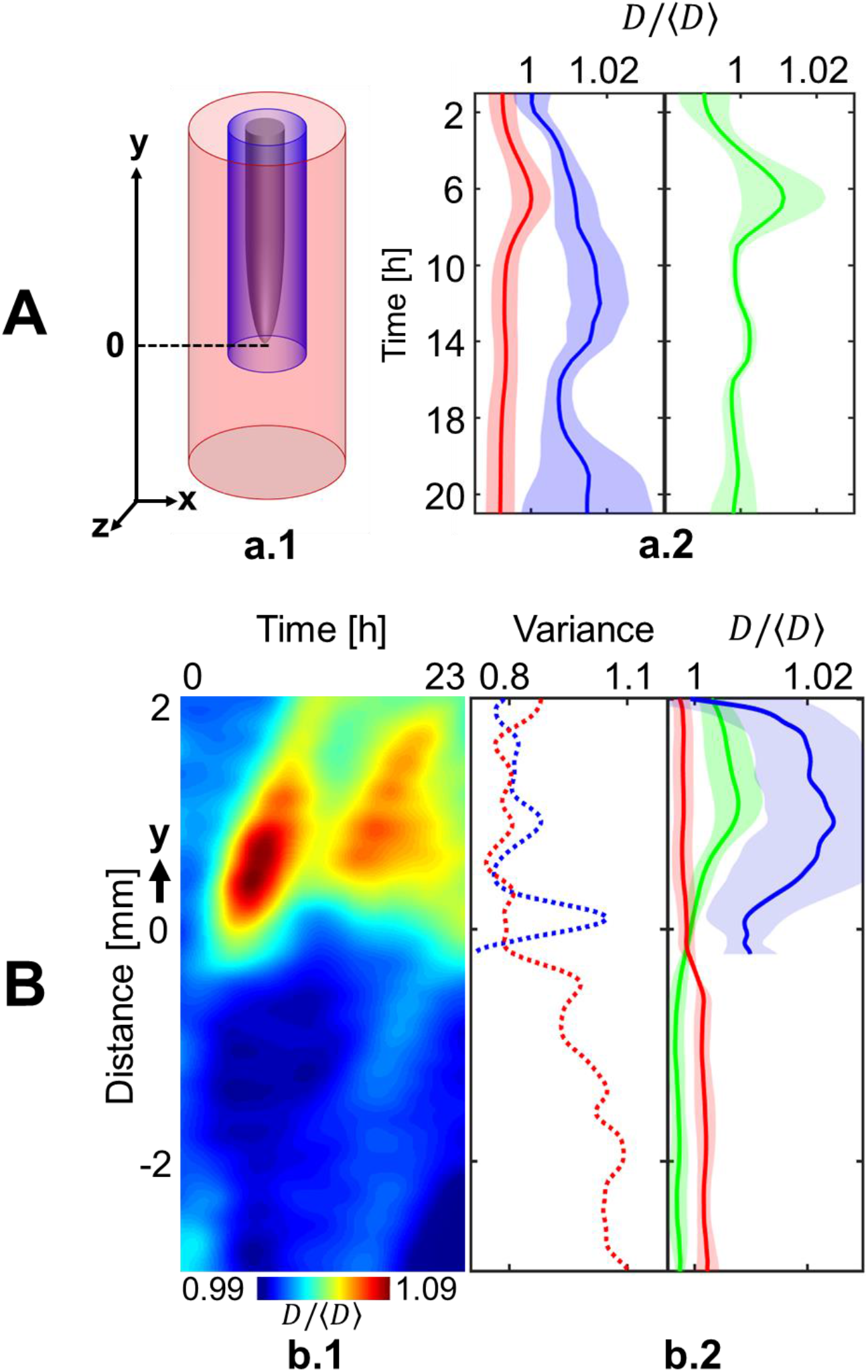
Dynamics of the colonisation of lettuce roots by *Bacillus subtilis*. (A) The colonisation is marked by an increase in bacterial cell density in the soil further away from the root (red) within the first 6 hours following inoculation, and a subsequent densification on or closer to the root surface (blue) until a steady state is reached between 10 and 14 hours following inoculation. The total quantity of bacteria in the system (green) reaches a steady state after 6h, which indicates migration not growth may play a role in the later stages of the colonisation of the root. (B) Intense activity at the root tip may precede colonisation near or on the root surface. (Left) The diagram of the colonisation kinematics shows how cell density changes both with time and as a function of the position along one root. The diagram shows that densification of bacterial cell population is discrete (here two pulses 7 hours apart are recorded at approximately 1 mm from the tip) and likely results from the attachment of bacteria on non-growing tissue since diagonal patterns indicate constant increase in the distance from the root tip. (Middle) Overall, the distribution of bacterial cell density along the root (solid blue line) confirms bacterial cell density concentrates in the basal region of the root (>1mm from the root tip). (Right) On the contrary, the most intense temporal variations in bacterial cell density are observed near the root tip (dashed blue line). The activity of bacterial cells in the bulk soil (red) confirmed also enhanced activity at the tip of the root, with both the density (solid red line) and the variance (dashed red line) showing a maximum in the region near the root tip. Data shown as mean ± SE, n=6.

To understand whether hot spots of bacteria move along the root, diagrams of the colonisation kinematics of individual roots were obtained (Figure 5 B, left). Bacterial hotspots (high bacterial cell density) appeared as diagonal stripes in the space-time domain, which showed bacterial hotspots were mostly immobile. Hot spots are formed, therefore, from bacteria converging towards sites of attachment on the root surface or on surrounding soil particles.

The system did not allow the tracking of individual cells across the entire soil volume. To understand the movements of cells preceding attachment, we studied instead the variance in bacterial cell density (equation 4) along the root and we could link the variance with regions the cell density. As described earlier, the bacterial cell density was found to be larger in the basal and more mature part of the root, at a distance starting approximately 1 mm from the root tip (Figure 5 B solid blue line). In contrast, bacteria from the bulk soil concentrated in the opposite direction, at a distance larger than 500 μm from the root tip. The variance of bacterial cell density revealed the sites of high cell mobility and identified regions of soil where bacterial mobility was most intense. Large variances were recorded close to the root tip (Figure 5 B, dashed blue), and in the soil in front of the root tip (Figure 5 B, dashed red). This indicates the root cap may be a point of “first contact” for bacteria, with attachment and colonisation occurring at a later stage and on the elongation zone of the root.

## Discussion

Performing live microscopy on plant-soil-atmosphere systems is challenging because of the necessity to maintain suitable conditions to grow plants and microorganisms, both in soil and within the confined space of a microscope. Many of these constraints are inherent to the observation and imaging of living organisms (Udan et al., 2014; Keller et al., 2008). As a result of these challenges, and in spite of the importance of imaging organisms *in situ*, attempts to observe environmental interactions using live microscopy in solid structured media have remained extremely limited. To date, the vast majority of the understanding of root-microbe dynamics is based on experiments run in hydroponic or agar cultures (Massalha et al., 2017; Aufrecht et al., 2018; Carroll et al., 2020; Noirot-Gros et al., 2020) which bear little resemblance to reality, and by extraction where scales cannot be assessed properly.

At the other end of the spectrum, environmental or ecophysiological studies are limited by technologies for direct *in situ* observation of biological processes. Recent developments in neutron and X-ray tomography (Tötzke et al., 2019; Morris et al., 2017) have revealed the complex nature of the interactions between root systems and soils and the diversity of their responses to each other (Koebernick et al., 2019). However, the techniques available are slow and detection of microorganisms is limited or non-existent. Other techniques such as laser ablation tomography (Vanhees et al., 2020) promise fast quantification of biological structures of roots within undisturbed soil cores, but in a destructive manner.

Here, we have developed microscopy technologies that help bridge the gap between environmental and biological sciences. The technique combines the use of a transparent soil which allows control of water, light and nutrient supply within mesocosms and the circulation of refractive index matching liquid to image an entire plant and its surrounding environment. Using the system, root-soil-bacteria interactions were observed *in situ*, and we have gathered evidence that soil microstructure affects bacterial behaviour.

Around root tips, where bacteria were more mobile, the occupation of the pore space was evenly distributed. However, when bacteria established on or in the vicinity of the root, high bacterial cell densities were observed primarily in smaller pores, confirming experiments made on fixed samples (Vanek and Thies, 2016; Raynaud and Nunan, 2014). Soil heterogeneity is known to limit microbial mobility (Young et al., 2008), and we found that it may cause bacteria to grow and appear in pulses, targeting root surfaces as a group and forming patches on the root surface. The study therefore provides first experimental evidence on how soil structure can modulate the dynamic behaviour of bacteria around growing roots.

Observations also hinted at the possibility that complex movements of cells occur before attachment on and colonisation of the root surface. As described earlier, *Bacillus subtilis* move chemotactically towards root surfaces (Allard-Massicotte et al., 2016), and attachment has been observed 1 mm from the root apex, in a region that corresponds well to the end of the elongation zone (Massalha et al., 2017). However, the increased sensitivity and ability to monitor microbial dynamics allowed us to identify other regions of importance to the microbial colonisation process. A peak in bacterial mobility was consistently observed at the root apex. This peak of activity differed from the accumulation of cells in the root tip seen in other bacterial species (Watt et al., 2006) because high mobility was associated with low bacterial cell density. This activity could be linked to attraction and interaction with specific cell types, e.g. border cells, and mucilage released by the root cap (Vicré et al., 2005).

This study demonstrates the ability of live microscopy to observe plant and micro-organisms within their complex environments. Continued efforts are now critical to integrate emerging optics technologies and deliver a first generation of environmental microscopes. The potential of this development to promote our understanding of the biology of this critical environment is enormous.

## Material and Methods

### Fabrication of chambers

Plants and bacteria were grown in mesocosm chambers holding soil, water and nutrients. Chambers consisted of glass slides (76×26×1 mm^3^, VWR, UK) and polydimethylsiloxane (PDMS, SYLGARD^TM^ 184, Sigma-Aldrich, UK). PDMS with a 3×3 mm^2^ cross sections was used to seal the glass slides and for flexible supply of gas and fluids using syringes. Therefore, the chamber obtained had a volume of 4560 mm^3^. Nutrients and index matching liquid were infiltrated into the soil using two Ismatec Reglo peristaltic pumps (Cole-Parmer, Wertheim, Germany) when required.

### Multispectral Light Sheet Fluorescence Microscopy (LSFM) for whole plant-environment imaging

The Gaussian beam from a four-channel laser source (488, 514, 561 and 633 nm, VersaLase^TM^, Laser 2000, UK) was expanded to 2.6 mm in diameter (FWHW) and split evenly into two illumination arms (Supplementary Information 1&2). The homogeneous light sheet was generated using two Powell lenses (10° fan angle, LOCP-8.9R10-2.0, Laser Line Optics, Canada) and two cylindrical lenses (100 mm focal length, LJ1567RM-A, Thorlabs, UK). The beam thickness has a full width half maximum of 50 μm, with a measured Rayleigh range of 1.7 mm. The image was projected through a 2× NA=0.055 or 5× NA=0.14 long working distance objective (Mitutoyo Plan Apo Infinity Corrected), a fluorescence emission filter changer (Four-Position Slider, ELL9, Thorlabs, UK) and a tube lens (TTL200-A, Thorlabs, UK) to a scientific camera (CMOS Camera, C11440-22CU, Hamamatsu, UK). A 3-axis translation stage was used for acquisition of large volumetric data. It consisted of two DC motor linear stages (M-VP-25XA, MKS Newport, UK) for horizontal displacement and a stepper motor linear stage (LNR50S/M, Thorlabs, UK) for vertical displacement. Fluorescence and scattering signals were acquired serially. With a 2× objective 7×60×35 mm^3^ can be imaged from image-stacks of 200 slices (in step of 50 um) at 10 vertical positions obtained in steps of 4 mm. Since the chambers were 3 mm thick, the total volume of the sample imaged was 3600 mm^3^. Illumination using 633 nm was used to collect scattering signals generated by the plant. All other signals were used to collect fluorescence signals with a band pass (520nm, 36nm Edmund Optics, UK) or a series of long pass (550nm, 600nm, and 650nm, Thorlabs, UK) emission filters. The chamber was attached to the 3-axis stage by a custom made holder. The holder was attached to a manual rotation stage (MSRP01/M, Thorlabs, UK), and samples were positioned and translated along an axis forming an angle of approximately 45 degrees with the illumination and detection axes. The sample was placed in an acrylic tank filled with approximately 10% sugar solution (RI=1.34).

### Environmental control

Transparent soils are produced from granular substrates whose particles are made from transparent materials that have a refractive index approaching that of water (1.333). Nafion^®^ in the form of pellets was used to generate the transparent soil particles (4mm × 3 mm, Ion Power Inc., USA). The particles were fractured to a size similar to those found in sandy soils (0.25 to 1.25 mm) using a freezer mill (6850 Freezer/Mill, SPEX CertiPrep, UK) and a series of sieves. pH and mineral ion concentration on the surface of the particles was then obtained by a series of chemical processes described earlier (Downie et al., 2012). Percoll (colloid suspension, GE Healthcare, UK) was infiltrated into the soil to match the RI of the Nafion^®^ particles before imaging. Plants grew under the illumination generated by a LED light panel composed of red and blue light with 3:1 ratio and producing photosynthetic photon Flux of 240 μmol s^-1^. The arrangement of the LED was designed to fit the sample holder and rotary stage. Water from the tank was circulated with miniature water pumps (480-122, RS Components, UK) through a Peltier device (RS 693-7080, Components, UK) with the temperature controlled using a TLK33 controller (Ascon Tecnologic, Italy). During the experiments the temperature was set to 20 °C.

### Plant and bacterial culture

Seeds of lettuce (*Lactuca sativa* all “year round”, Sutton Seeds, UK) were surface sterilized by washing in 10% bleach for 15 minutes followed by thorough rinsing with sterile dH_2_O before overnight germination. A single seedling (with approximately 2 mm root length) was then transferred into each mesocosm filled with transparent soil saturated in Murashige and Skoog Basal Medium (MS, Sigma Aldrich, UK) and stained with sulforrhodamineB. *Bacillus subtilis* NCIB 3610 GFP labelled strain (NRS1473(Hobley et al., 2013)) was grown in MSgg medium (5 mM potassium phosphate and 100 mM MOPs adjusted to pH 7.0 and then supplemented with 2 mM MgCl_2_, 700 μM CaCl_2_, 50 μM MnCl_2_, 50 μM FeCl_3_, 1 μM ZnCl_2_, 2 μM thiamine, 0.5% glycerol (v/v), and 0.5% (w/v) glutamate) for 28 hours at 18 °C, whilst shaking at 200rpm. After incubation, the MSgg solution was replaced with MS to remove any carbon contained in the bacterial solution. Based on the OD_600_ absorbance of the bacterial suspension in half MS and the known correlation with CFU for this strain, approximately 2.0 × 10^6^ CFU were inoculated onto a sterile filter disc. The inoculated disk was then inserted into the mesocosm, on the surface of the transparent soil, next to the plant seedling. Mesocosms with a lettuce seedling in each, were inoculated and subsequently incubated at 21°C for 20 to 24 hours before introducing the Percoll solution for index matching and imaging. Images capture was initiated in the morning and collected the following day. In total, 8 individual mesocosms were inoculated with bacterial suspension and 3 individual mesocosms grew without inoculation of bacterial suspensions for assessing the effect of bacteria on root growth.

### Software control

The environmental microscope was controlled through custom made LabVIEW software (National Instrument, Austin, Texas, USA). A single-board microcontroller (Arduino Mega 2560, RelChron Ltd, UK) controlled of the laser output through RS232 external triggers. The growth light was powered by a DC power supply controlled by a USB-RLY08 relay (Devantech Limited, UK).

### Image acquisition and processing

Data processing was tailored to requirements for quantification of root and particle geometry and quantification of bacterial cell density (Supplementary Information 3). Fluorescence and scattering signals were processed using affine transformation and nearest neighbour interpolation to correct for the angle of the scan (45°) used. Volume data was subsequently processed by the Lucy-Richardson deconvolution method with a light-sheet point spread function (Becker et al., 2019). Overlapping regions were fused using a Laplace pyramid blend (Gonzalez and Woods, 1990; Burt and Adelson, 1983). As soil is a textured material, it contains periodic (particles) that can be used to infer flat field corrections. The correction was based on a weight matrix computed from the mode value of the pixel intensity computed from neighbouring pixels of an entire dataset, and modification of the image intensity applied using Laplace pyramids method.

Image segmentation was used to extract the shape, structure and spatial distribution of roots, bacteria and soil particles (Supplementary Information 4). The morphology of the root was obtained using a region growing algorithm and the resulting binary data was used to produce distance maps and to calculate the position of bacteria relative to the root surface. Only minor changes to the threshold values and position of the seeds were needed to account for variations in scattering intensity and movements of the root. Because sulforrhodamine-B attaches only superficially to the soil particle, the signal was not sufficient to segregate the pore space from the core of the particle. Therefore, we overplayed the inverse of the GFP signal to improve the segmentation of the pore space, and a manual threshold followed by morphological operators (dilation and erosion) generated binary images of the pore space. The pore size was calculated using the local thickness metric (Supplementary Information 4).

Image processing methods were programmed using MATLAB^®^ using the Image Processing Toolbox (MathWorks, USA). Segmentation and extraction of geometrical features were performed using MeVisLab^©^ (MeVis Medical Solutions AG, Germany). All software is freely available from http://archiroot.org.uk/.

### Calibration of fluorescence signal

Dense bacterial suspensions were prepared in Percoll and measured by OD_600_. Suspensions prepared at 12 different Optical Density values (OD) in the range 1.2 10^-3^ to 3.0 were obtained by multiple dilutions. One ml of each bacterial suspension was transferred into mesocosm chambers and stained soil particles were added to the chamber to adjust the focus of the environmental microscope. A full scan was acquired 2 mm above the particles (in steps of 50 μm and at two z levels 4 mm apart). Pixel intensity data was then correlated against OD values, and OD values were correlated to CFU counts. The estimation of bacterial cell density from image data was based on the combination of both correlations (Supplementary Information 5).

### Indicators for bacterial activity

Different indicators were used to map bacterial activity in the pore space. Bacterial cell density was estimated from the intensity of the green fluorescence signal. For all analyses, a pixel at a given time point is associated with three variables, pore-size, distance from the root surface, and distance from the tip of the root. Pixels were subsequently classified into groups (*R_k_*) related to their relative position along the root, to their position perpendicular to the root or the size of the pore they are located in.

For each group of pixels *R_k_* at time *t*, the bacterial cell density (ml^−1^mm^−3^) is defined as the estimated number of bacterial cell (CFU) in a unit soil volume expressed per unit volume of root (Carroll et al., 2020),

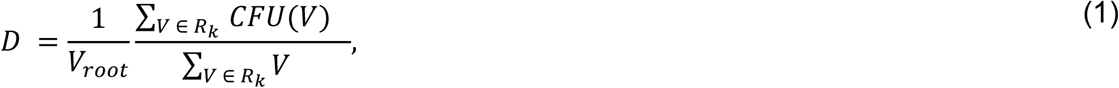

where V_*root*_ is the volume of the root. To allow for comparisons between experiments we calculated the relative bacterial density at time t,

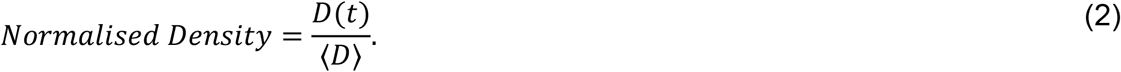

The expected value of the bacterial cell density ⟨*D*⟩ is calculated across the entire experiment of duration *T*,

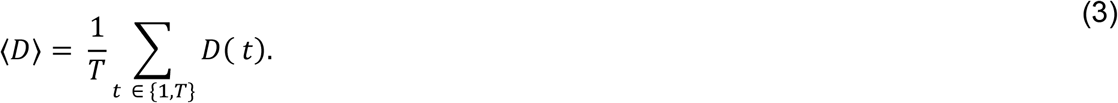

The mobility of bacteria was quantified as the variations in bacterial density at a given location in soil. This was calculated using the temporal variance of the bacterial cell density,

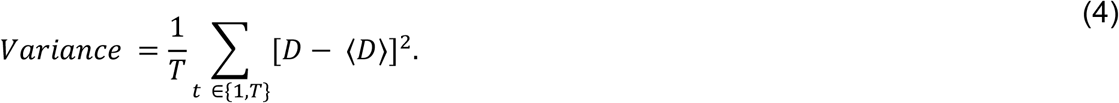

The first 5 time points were discarded from the computations to allow sufficient soil volume in the mature region of the root. Rhizosphere pixels were defined as pixels for which distance from the root surface is less than 0.2 mm. Bulk soil pixels are defined as pixels which distance is more than 0.2 mm from the root surface. Pixels are associated with the root tip when their projection along the root is less than 2 mm away from the tip, and pixels associated with the base of the root are more than 2 mm away from the tip.

## Acknowledgments

We thank Alberto Lora de la Mata and Ana Garcia Verdugo Zuil for helping with the preparation of chambers and transparent soil. We are grateful to Jacqueline Marshall and Maud Darsonval for assistance with microbiology work. We thank Ian McNaughton for assistance with the design of electronic components. The James Hutton Institute & SRUC were supported by funds from the Rural and Environment Science and Analytical Services Division of the Scottish Government. Work in the NSW laboratory is funded by the Biotechnology and Biological Science Research Council (BBSRC) [BB/P001335/1, BB/R012415/1].

