## Supplementary Methods for "Plant-environment microscopy tracks interactions of Bacillus subtilis with plant roots across the entire rhizosphere"

### 1 - Microscope setup

A Gaussian beam from a four channel laser source (wavelength: 488, 514, 561 and 633 nm, VersaLase™, Laser 2000, UK) was expanded to a diameter of 2.6 mm at the Full Width at Half Maximum (FWHM) and split evenly into two illumination arms (Figure 1). A homogeneous light sheet was generated by two optical Powell lenses (10° fan angle, LOCP-8.9R10-2.0, Laser Line Optics, Canada) and two cylindrical lenses (LJ1567RM-A, Thorlabs, UK) in each illumination arm. The selected cross-section was projected to a scientific camera (CMOS Camera, C11440-22CU, Hamamatsu, UK) through a 2X/0.055 or 5X/0.14 long working distance objective (Mitutoyo, Plan Apo Infinity Corrected Objectives, Edmund Optics, UK), a filter changer (ELL9, Thorlabs, UK) and a tube lens (TTL200-A, Thorlabs, UK).

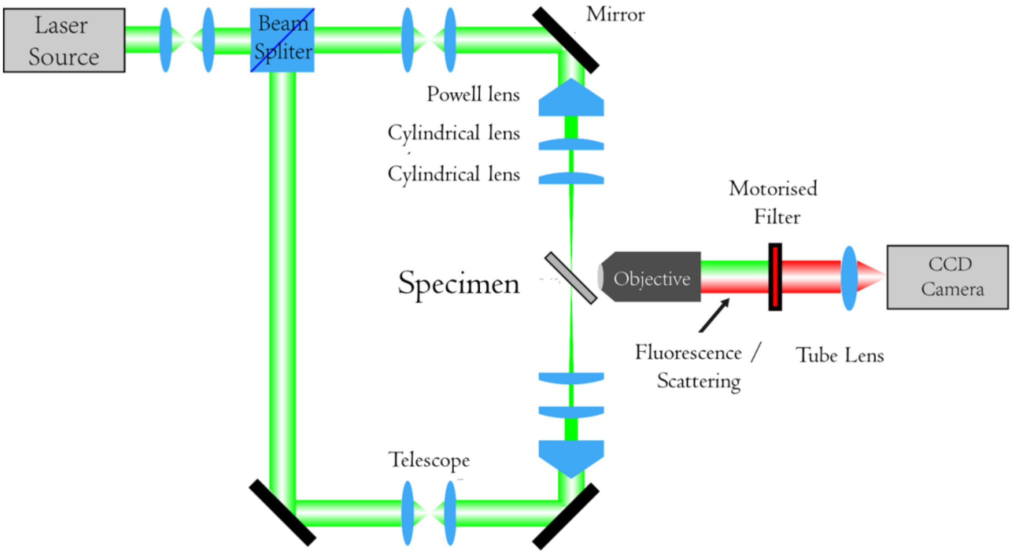


Figure 1. Design of Light Sheet Fluorescence Microscope for whole plant-environment imaging.

A 3-axis translation stage was assembled for scanning the light sheet through the whole sample. It consisted of two single-axis DC motor linear stages (M-VP-25XA, MKS Newport, UK) for displacement in the horizontal plane and a stepper motor linear stage (LNR50S/M, Thorlabs, UK) for displacement along the vertical axis. The chamber was fixed to the stage by a custom-made clip holder, which itself was attached to a manual rotation stage (MSRP01/M, Thorlabs, UK) for positioning of the chamber at approximately 45 degrees to the illumination and imaging axes.

The chamber was immersed in an acrylic tank (Figure 2) filled with 10% sugar solution (RI=1.34). The growth light was attached above the chamber holder and connected to a DC power supply controlled by a USB-RLY08 relay (Devantech Limited, UK). A pair of ismatec reglo peristaltic pumps (Cole-Parmer, Wertheim, Germany) was used to circulate the fluids to the chamber using Teflon® tubes (1.2

mm for suction and 0.8 mm for injection). Needles were attached to the end of the Teflon® tubes and inserted through the PDMS spacers of the mesocosm chambers.

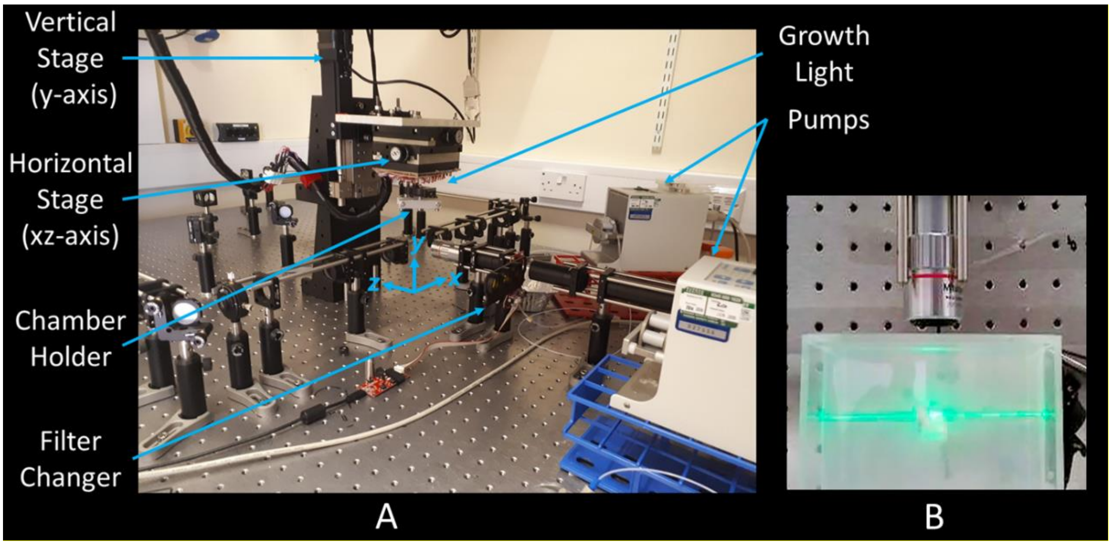


Figure 2. Annotated photograph of the light sheet microscope. (A) Overall view of the set up. (B) Close up above view the tank used for refractive index matching and temperature control located under the stage (indicated by the xyz axes in A).

### 2 - Characterisation of microscope

The light-sheet was measured using a CMOS camera (DCC1545M, Thorlabs, UK). The camera was equipped with a neutral-density filter and was mounted on a LNR50S/M linear stage (Thorlabs, UK) with the sensor placed perpendicularly to the light-sheet. The laser power output was set to 1 mW. The stage translated the camera along the light sheet with a step size of 20 µm. The image data was collected to reconstruct the 3D light intensity distribution of the light-sheet (Figure 3). The measured beam waist of the light-sheet was 50 µm (at FWHM), the Rayleigh range was approximately 3 mm, and the height of the beam was 7.6 mm (at FWHM).

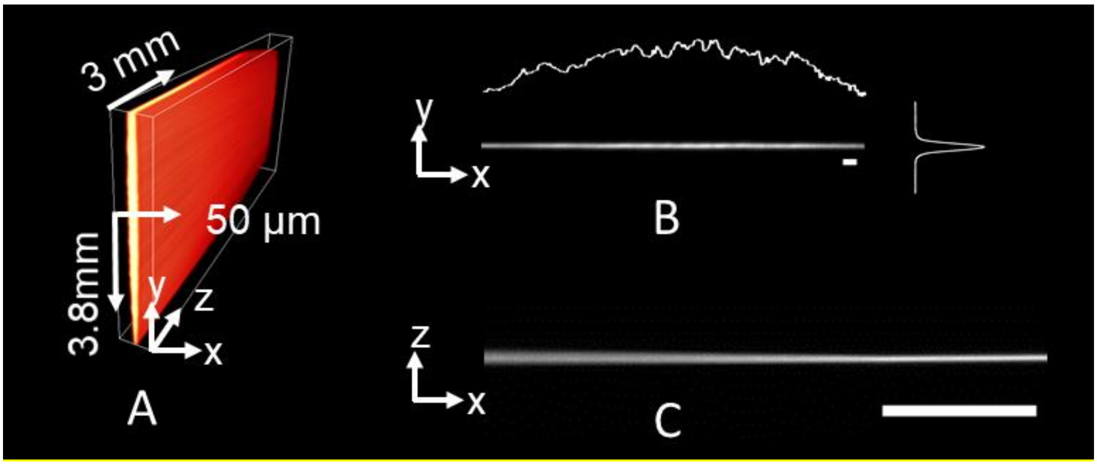

Figure 3. Measurment of the light sheet properties. (A) 3D light intensity distribution of the light-sheet obtained from a camera capturing the cross section of the light sheet along the axis of propagation of light. (B) Beam profile at the waist. The scale bar is 100 µm. (C) Beam profile along the direction of propogation. The scale bar is 2 mm.

Microspheres of 10 µm diameter with green-yellow fluorescence (505/515 nm, ThermoFisher, UK) were used for the characterisation of the resolution of the microscope. The microspheres were immobilised in 1% agar (Sigma-Aldrich, UK) in 1×1×5 cm spectrophotometer cuvettes (58017-875, VWR, UK). The sample was mounted on the stage of the microscope and the lateral resolution of the microscope was determined using the 5X/0.14 objective. The shortest distance measureable between adjacent microspheres was determined to estimate the lateral resolution (Figure 4 A&B). The shortest distance observed was 13 µm, but the actual resolution is likely less than 10 µm because the microsphere diameter was significantly larger than the resolution of the objective (around 2 µm). Microsphere samples were also used to align the light-sheets. Misalignments we observed using the 5X objective (Mitutoyo) and adjustments were performed until no difference were observed between single and dual illumination.

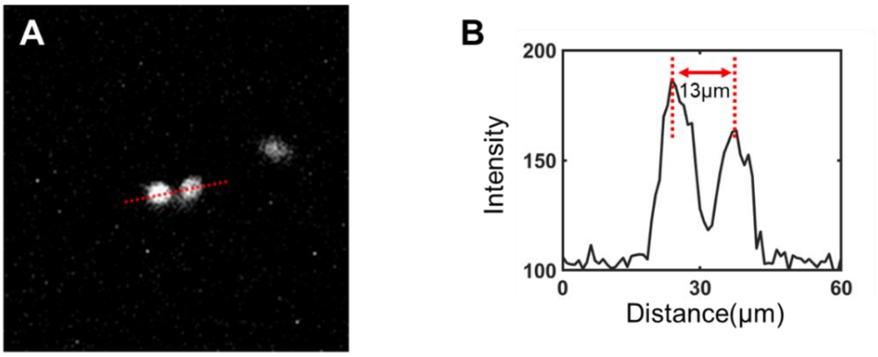


Figure 4. Estimation of lateral resolution. (A) Two adjacent microspheres. (B) Image intensity profile along the line joining the two particles (red dashed line in A).

#### 3 - Processing of image data

This section describes the development of a pipeline for processing the data collected by our light sheet microscope. The processes include affine transformation to correct for the scanning angle, 3D deconvolution, flat field correction, fusion and image stitching. The processing of the data is essential to all subsequent quantitative analyses of roots, soil and bacteria presented in the following sections. Unless specified otherwise, all processes are implemented in custom made software using the MATLAB® (MathWorks, USA) programming language (Figure 5).

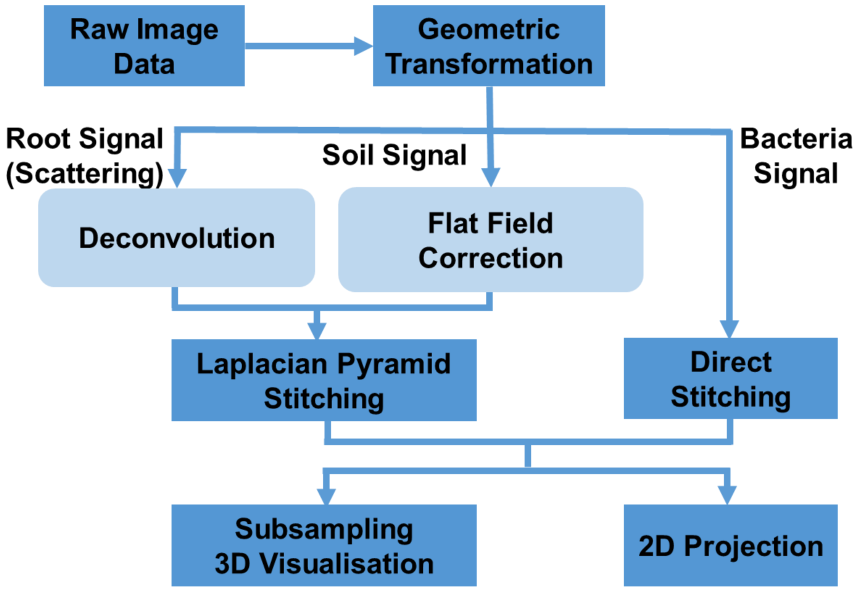


Figure 5. Image processing pipeline. Geometric corrections are needed to correct for the scanning angle. Here the correction is an affine transformation to restore distortion due to scanning at a 45 degree angle with the imaging arm. Deconvolution is applied to the root signal (scattering) to improve contrast from the noise generated by the elastic scattering. Statistical estimation of flat field correction is subsequently obtained. Finally, a stitching method is developed. It combines multilevel blending and flat field corrections estimated at the previous step. The data assembled can then be downscaled for computationally intensive tasks such as visualisation or image analysis. Alternatively, projection algorithms can be used for fast 3D visualisation a maximum resolution. Dark blue colours represent tasks performed independently of the type of signal collected. Light blue colours represent tasks that are dependent on the type of signal collected.

#### Geometric transformation

The imaging chamber is translated along an axis that makes a 45° angle with the illumination and imaging axes. Therefore, it is necessary to apply a first geometric transformation to map all pixel values acquired by the camera into the global orthonormnal coordinate system, defined by the axis of the illumination axis, the axis of imaging, and the vertical axis (Figure 6).

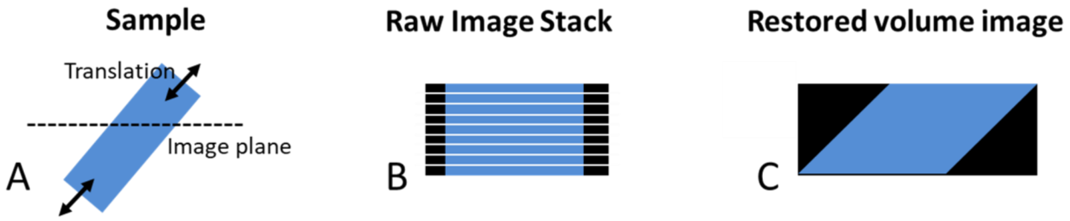


Figure 6. Geometric transformation of volume data. (A) The sample is moved at a 45° angle to the imaging axis. (B) When images obtained are stacked into a volume

dataset, the shape of the sample (in blue) is distorted. (C) Geometric restoration corrected for such distorion to build a volume image with dimensions that coincide with those of the sample.

The original position of a given pixel in the global coordinate system is obtained by an affine transformation whose matrix is expressed as a function of the size of the step of the translation stage $d_s$, the slice number $i_s$, and the angle at which the translation is made $\theta$, e.g. 45 degrees. The transformation is defined as

$$\begin{bmatrix} x' \\ y' \\ z' \\ 1 \end{bmatrix} = \begin{bmatrix} ds\,sin(\theta) & 0 & 0 & 0 \\ 0 & 1 & 0 & 0 \\ ds\,cos(\theta) & 0 & 1 & 0 \\ 0 & 0 & 0 & 1 \end{bmatrix} \begin{bmatrix} i_s \\ y \\ z \\ 1 \end{bmatrix}. \qquad (1)$$

For a 5X/0.14 objective the step size *ds* is 30 µm and for 2X/0.055 the step size is 50 µm. The size of the horizontal step is generally larger than the resolution of the camera. Rescaling of the data can also be included at this stage to obtain voxel of identical size in the three dimensions. The affine transformation was performed with nearest neighbour interpolation.

**Deconvolution**

We have tested different models for point spread function (PSF). The PSFs tested included the Born & Wolf PSF model from the imageJ plugin PSF Generator [1], Pankajakshan's model [2] and Becker's light sheet microscope model [3]. Tests of the different PSF functions were first made on image data generated from microsphere test samples before application to live imaging data.

*Test data*

Test samples were made of 10 µm FluoSpheres™ fluorescence microspheres (505/515 nm, F13081, ThermoFisher, UK) and 1% agar. One µl microspheres were dispersed in 1 ml water and then further diluted by adding 4 ml of 1% melted agar solution. The microsphere agar suspension was shaken and then poured into acrylic spectrophotometer cuvettes (58017-875, VWR, UK) before cooling at room temperature. A final dilution ratio of 1:5000 was obtained. The immobilised beads were scanned horizontally along a distance of 300 µm with a step size of 3 µm. The step size was chosen to maintain the axial resolution to the same level as the theoretical resolution of the microscope objective. Both Lucy-Richardson algorithm and Wiener filter algorithms for deconvolutions were tested. Visual assessment of the results showed the Lucy-Richardson method performed marginally better than the Wiener method, and it was used for the remaining analyses.

*Point Spread Functions*

The image data collected from the microsphere test samples was used to estimate the

Point Spread Functions (PSF). The data showed strong distortion in microsphere geometry, largely due to mismatches between the light sheet and the focusing plane of the imaging optics (Figure 7 A). First, the PSF was modelled using the Born & Wolf model which corrects for scalar diffraction occurring when there is a mismatches between the position of the point source and the focus of the objective. This model can account for artefacts created by low Numerical Aperture (NA) objectives. The Born & Wolf model was best fitted with a misalignment of 7 µm (Figure 7 B). Pankajakshan's model is also a scalar diffraction limited model but with spherical aberration (Figure 7 C). A third point spread function was generated based on the Becker's light sheet microscopy model using parameters such as focusing distance, entering pupil and Numerical Aperture (NA) of the illumination arm (Figure 7 D). A model was also obtained using Pankajakshan's model confined by the measured thickness of the light sheet. In this model, we assumed a perfect alignment between the focus of the illumination and the imaging optics (Figure 7 E). The beam distribution near the target was first transformed to have a norm of 1, and then used to correct the Pankajakshan's PSF function. Finally, Figure 7 F shows the Pankajakshan's model confined by the light sheet function with misalignment (Figure 7 F). The best fit was obtained with a misalignment of 10 µm. The light sheet properties used to build the different models were obtained from direct measurements of the light sheet profile (Figure 7 G).

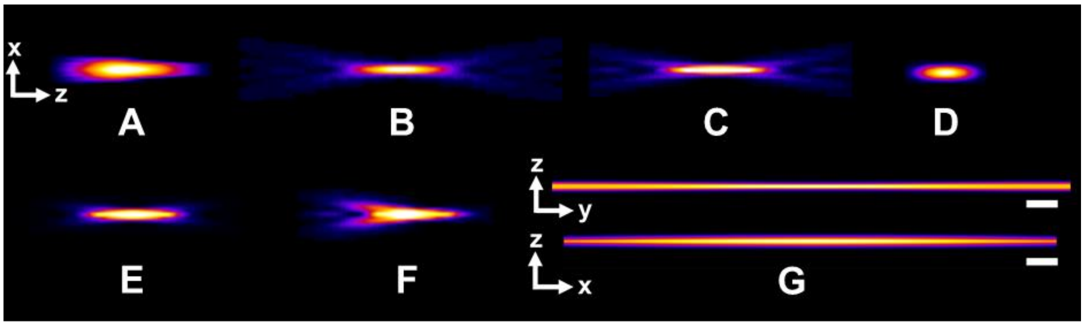


Figure 7. Point spread function (PSF) measured and estimated by various models. (A) point spread function produced experimentall by 10 µm bead. Models tested for the dconvolution of the data include Born & Wolf model (B), Pankajakshan's model (C), Becker's light sheet microscope model (D), Pankajakshan's model constrained by a light sheet (E), Pankajakshan's model confined by a light sheet with 10 µm misalignment (F). (G) Smooth illunimation beam profile used to generate models in (D-F). The scale bar is 1 mm in size.

*Deconvolution of volume data*

The point spread functions shown in Figure 7 were used to perform deconvolution of the 10 µm microsphere image data. Deconvolution results were assessed visually from the volume rendering of the deconvoluted dataset (Figure 8). Results showed that the PSF models of Born & Wolf (Figure 8 B) and Becker's light sheet microscope model (Figure 8 D) can restore the distored signal of the sphere back into a reasonable shape by comparison to the results obtained by the Pankajakshan's model (Figure 8 C). Pankajakshan's model confined by the measured thickness of the light sheet further improved the restoration of the original shape of the sphere (Figure 8 E). Addition of misalignment between the light sheet and the focal plane of the microscope objective (Figure 8 F) allowed complete recovery of the spherical shape of the particle.

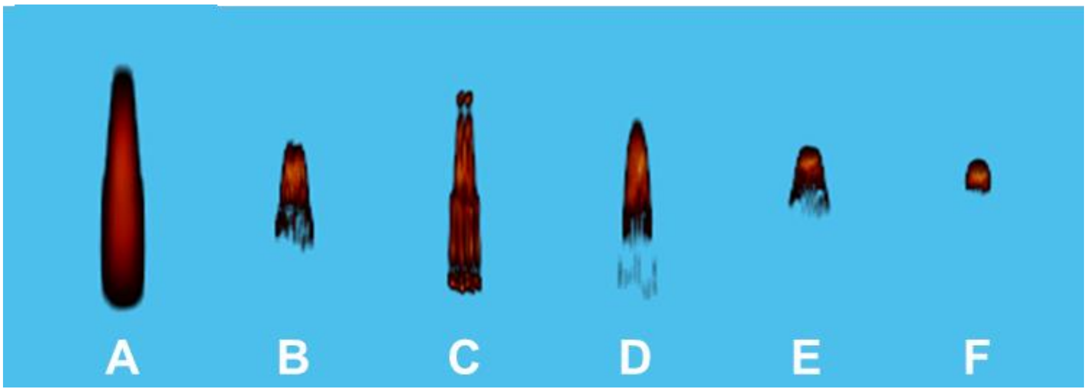


Figure 8. Volume redering of deconvoluted microsphere data. (A) 3D visualisation of a 10 µm bead without deconvolution. (B) Deconvolution with the Born & Wolf model. (C) Deconvolution with Pankajakshan's model. (D) Deconvolution with Becker's light sheet microscope model. (E) Deconvolution with Pankajakshan's model confined by the thickness of the light sheet. (F) Deconvolution with Pankajakshan's model confined by the thickness of the light sheet and with 10 µm misalignment.

In a second step, deconvolution methods were applied to image data from roots growing in transparent soil. Results showed the type of point spread function used for deconvolution affected the quality of the recovery of the root structure. Methods based on the Born & Wolf model (Figure 9 B) and confined Pankajakshan's model (Figure 9 C) performed surprisingly well in comparison to the confined Pankajakshan's model including misalignment (Figure 9 D). In these models, cell files could be observed more clearly than on the original data (Figure 9 A) and images showed enhanced contrast as a result of deconvolution.

Results in Figures 8 & 9 established that the deconvolution significantly improves the axial resolution of the microscope. The best results are obtained when the optical properties of the imaging system are known with sufficient precision. In practice, precise measurement of the alignment of the light sheet with the focal plane is difficult to obtain due to the multiple air-solid-liquid interfaces the light must cross before reaching the camera sensor. Additional unknowns arise due to the heterogeneity of the soil medium, the angle of the sample, or small sample movements that occur during mounting on the microscope. Deconvolution with incorrect misalignment is detrimental to image quality, and simpler methods such as the Born & Wolf model or Pankajakshan's model confined model may thus prove to be more suitable for routine use of the microscope. In all cases however, all deconvolution methods removed out of focus light from the image and significantly improved subsequent processing and quantitative analysis performed on the data.

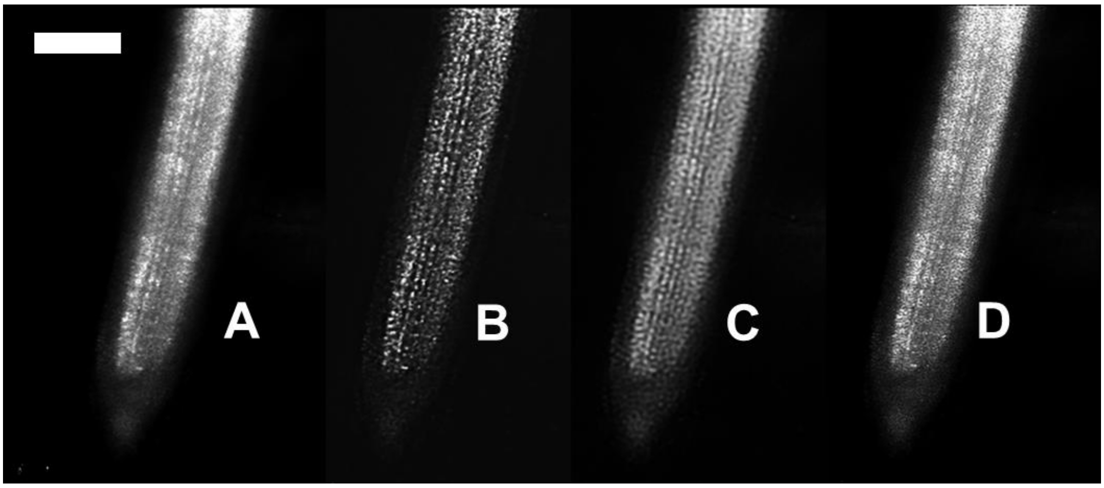


Figure 9. Deconvolution of lettuce root grown in Nafion® soil. (A) Average projection image of unprocessed data. (B) Average projection image of data deconvoluted with Born & Wolf model. (C) Average projection image of data deconvoluted with Pankajakshan's model confined by measured light sheet. (D) Average projection image of data deconvoluted with Pankajakshan's model confined by measured light sheet and with 10 µm misalignment. Scale bar 500 µm.

**Flat field correction**

Inhomogeneous contrast in raw images is a common issue in microscopy and imaging. It usually results from uneven illumination, optical distortions and intrinsic noise from the imaging sensor [4]. The process to remove this effect is known as flat-field correction. Flat-field correction is typically achieved using the formula [5,6]

$$C(x,y) = \frac{R(x,y) - D(x,y)}{F(x,y) - D(x,y)} \times m\,. \tag{2}$$

Two types of image data $D(x,y)$ and $F(x,y)$ are required for calibration before transforming the raw image $R(x,y)$ into the corrected image $C(x,y)$. $D$ is usually the dark field image captured with the same exposure time as those used during experiments but in the total absence of light. $F$ is the image acquired under the same illumination as the raw image but without samples. $m$ is taken as the average pixel intensity of $F$. $(R – D)$ corrects for the electronic offset of the sensor and $m/(F – D)$ restores the remaining optical distortions.

Flat-field correction is particularly important for the stitching of large dataset because variations in image intensity arise due to both the illumination and detection part of the acquisition process. The sample was mounted at an angle and placed in a liquid container with refractive index matching solution, and the imaging optics induce vignetting-like effects that are exacerbated when imaging through multiple interfaces (air – acrylic – RI solution – glass – soil). The flatness of the focal plane can also be degraded when using air objectives.

Standard flat field correction (Equation 2) cannot be easily applied to such complex imaging system. Obtaining data for performing flat-field correction in a light sheet set-up can be difficult to obtain without a target, since illumination is perpendicular to the imaging arm. Also the acquisition of adequate calibration data would significantly increase the time and complexity of experiments. Here, a method was developed to estimate flat field corrections from the raw image data itself. This is possible because the microscope was used in time lapse experiments which delivers data in large quantity from which flat field correction can be inferred. Secondly, the soil signal is particularly rich in information and covers the entire field of view.

The flat field correction developed is based on the following correction

$$C(x,y) = \frac{R(x,y)}{W(x,y)}. \tag{3}$$

The method generates weight factors $W(x,y)$ from the raw data using the probability density distribution $p$ of the pixel intensity $I$, determined at a given position $x, y$

$$p(I|x,y) = \frac{n_I(x,y)}{n}. \tag{4}$$

$n_I(x,y)$ is the number of pixels at position $(x,y)$ with intensity $I$ and $n$ is the number of observations at position $(x,y)$ . We then define the correction as a function of the argument maximum of the probability density distribution function,

$$W(x,y) \propto \frac{1}{arg\ max\big(p(I|x,y)\big)_I}, \tag{5}$$

so that the corrected image data has a conserved arg max of the pixel intensity. The algorithm processing the data first defines a box kernel with a size of 150 × 150 × 3900 μm (Figure 10). For each pixel on the image data, the histogram of the pixel intensity is computed from the neighbourhood generated by the box kernel. The mode of the pixel intensity is determined and the weight matrix $W(x,y)$ is computed.

The statistical flat-field correction method developed here benefits from the dark field and large volume of data generated by the field of view of the microscope. Also, the pixel intensity generally has a right-skewed distribution (Figure 10 A), where the number of background pixels is much larger than those of the foreground which increases the robustness of the method.

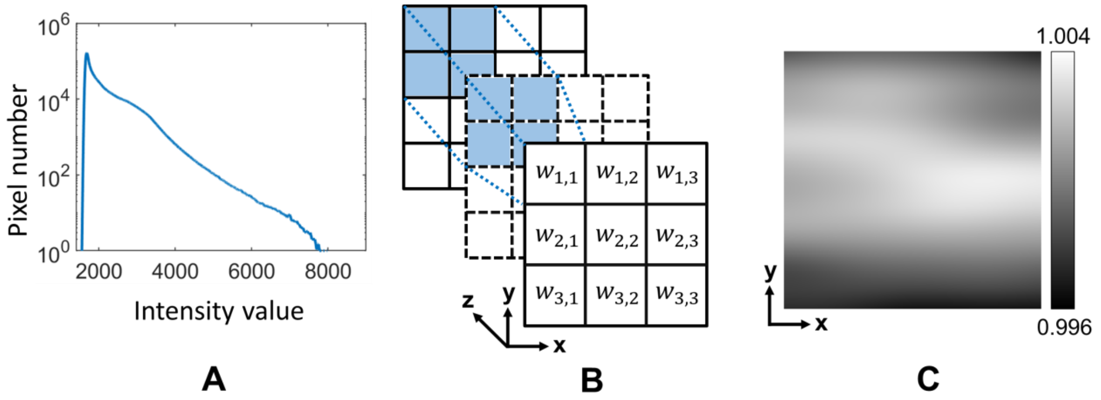


Figure 10. Flat field correction. (A) Histogram of a volume data dataset. (B) Weights for flat field corrections used the histogram obtained from neighbouring pixels (box kernel) accross an entire dataset. (C) Exampe of flat field correction obtained on a test dataset.

An alternative form of flat field correction was also developed (equation 6). In this case the weight value is obtained from the subtraction of the median from the mean of the unprocessed image intensity. This correction is similar to the skewness of a probability density distribution [7],

$$W(x,y) = \frac{mean(x,y) - median(x,y)}{mean[mean(x,y) - median(x,y)]}. \quad (6)$$

This approach is useful in the rare cases where the light sheet alignment had degraded with time.

The computed weight matrices usually showed moderate corrections (less than 1%) when light sheet alignment has been achieved with good precision (Figure 10). Flat field corrections were combined with multi-resolution blending which often improved the quality of the reconstructed (see below).

**Multiresolution pyramids stitching**

Volume datasets were assembled from different vertical positions by a process termed stitching. Stitching is the operation of using the overlapping domain of two images to create a combined image containing the most relevant information from either or both images. Stitching requires (1) identifying the overlapping domains and (2) determining the value of pixels in regions where the domains overlap and possibly in a neighbourhood that extends beyond the domain of overlap. The first operation maps both images in a common coordinate system using elementary transformations, e.g. translation, rotation or scaling, and is termed “registration”. Here, the overlap was determined by the vertical movement of the motorised stage of the microscope. The transformation was therefore a controlled vertical translation which creates an image overlap of 0.5 to 1 mm with a precision of 1 pixel or 0.1 µm, which is the resolution of the stage. The second step is termed “blending” and computes pixel values at the domains where images overlap, using a form of weighted average of pixel values. Here, the blending process was challenging because of the multiscale structures and the patterns present within images. We have used the Laplacian pyramid approach to address this issue.

Firstly, a raw image $I$ is convolved with a Gaussian kernel and then downscaled to an image half the original size. This elementary operation is termed $REDUCE$, and is iterated multiple times to generate a series of images $G(i)$,

$$G(i) = \begin{cases} I & , \quad i = 0 \\ REDUCE\big(G(i-1)\big), & 0 < i \leq N \end{cases} \tag{7}$$

The series of images $G(i)$ is termed a Gaussian pyramid (Figure 11).

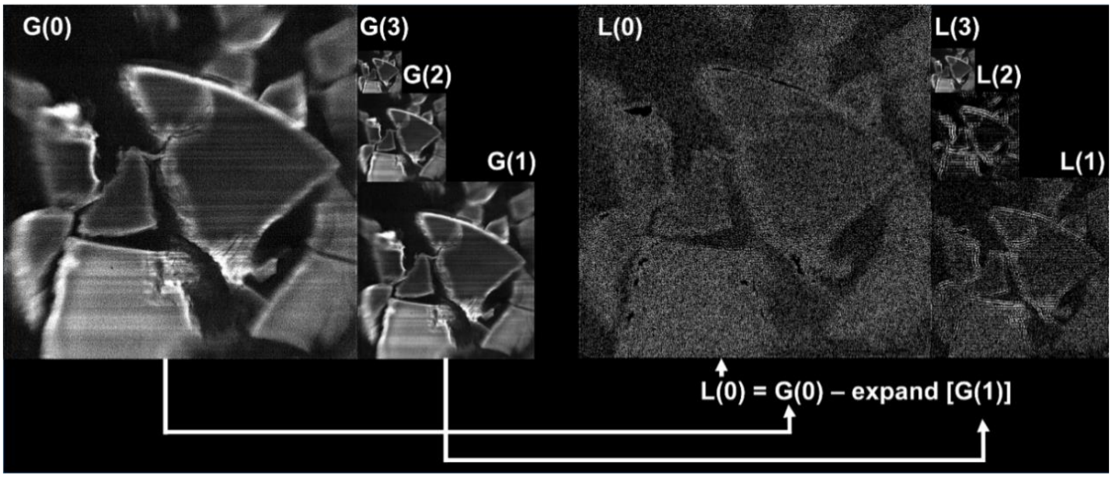


Figure 11. Construction of a Laplacian pyramid.

The Laplacian pyramid is then calculated from the Guassian pyramid,

$$L(i) = \begin{cases} G(i) - EXPAND(G(i+1)), & 0 \leq i < N \\ G(i), & i = N \end{cases} \tag{8}$$

In this case, a second operator termed $EXPAND$ is used to generate the pyramid. $EXPAND$ is the smoothed representation of the data at the lower scale and the Laplacian image $L(i)$ is obtained by subtracting the result of the i$^{th}$ $EXPAND$ from the i$^{th}$ Gaussian image. The process is initiated at the base of the Gaussian pyramid, $G(0)$, and iterated $N$ times to generate a second pyramid termed Laplacian pyramid (equation 8).

Because each level in the Gaussian pyramid is a smoothed representation of the lower scale, it can be interpreted as a low-pass filter and the Gaussian pyramid is a description of long range variations occurring at each scale within the image (Figure 11 left). On the contrary, each level in the Laplacian pyramid contains residual information after long range variations have been removed. It therefore acts as a high-pass filtering process describing small scale variation within at a given scale within an image (Figure 11 right). Various operations can then take place at each scale of the pyramid, including the blending of two images with overlapping domains or flat field corrections. Following operations at each level of the pyramid, an image can be reconstructed using the inverse relationship of equation 8, i.e. $g(i) = L(i) + EXPAND\ (g(i+1))$ .

*Method*

Since the image data generated by light sheet microscopy has both multiscale information and contains flat field artefacts the blending of overlapping values of pixel intensities must be achieved on a multiscale data structure. Also, to avoid vigneting effects, the blending must be done following flat field corrections. A mask ($\alpha(y)$ in equation 10) was also used to smooth the transition of two adjacent images. The weighting of the pixel intensity of blended images A and B of size $N$ in the $y$ axis and with an overlap of size $2e + 1$is

$$L_S(i) = \begin{cases} L_A(i) & ,for\ 1 \leq y < N - 2e - 1 \\ G_\alpha\ (i)L_A(i) +\ G_{1-\alpha}(i)L_B(i) & ,for\ N - 2e - 1 \leq y < N + 1 \\ L_B(i) & ,for\ N + 1 \leq y < 2N - 2e - 1 \end{cases} \quad (9)$$

The weighting function α is defined

$$\alpha = (k\ (y - N - e - 1))^2 \quad (10)$$

For blending using arithmetic average, α is assumed to take the value of 0.5. The algorithm used for stitching of light sheet microscopy images therefore follows the steps:

1. Compute Laplacian pyramid using equations 7-8;
2. Apply flat field correction to $G_A(i)$,$G_B(i)$, $L_A(i)$ and $L_B(i)$ following equation 3;
3. Blend $L_A(i)$,$L_B(i)$ using the following equation 9;
4. Reconstruct image using the reverse $EXPAND$ operator.

*Results*

The method developed was tested on the data produced from a small LED panel (smart phone) captured by the microscope equipped with the 2X objective. The weight matrix (equation 3) was generated by a minimum filter and applied to the raw image data and a 5-layer Laplacian pyramid was generated. The results were compared to a 5-layer Symlet wavelet representation.

The results of the test showed the method corrected successfully for flat field variations while preserving the multiscale structure of the image. The contrast of the foreground was significantly enhanced without enhancement of background information. Comparison with the original data (Figure 12 A) showed significant improvement of pixel intensity at the edge of the panel can be obtained using flat field correction. Following flat field correction applied to the Laplacian pyramid of the image, the image recovered an even brightness across the entire field (Figure 12 B). Similar results were obtained using the Symlet wavelet algorithm (Figure 12 C).

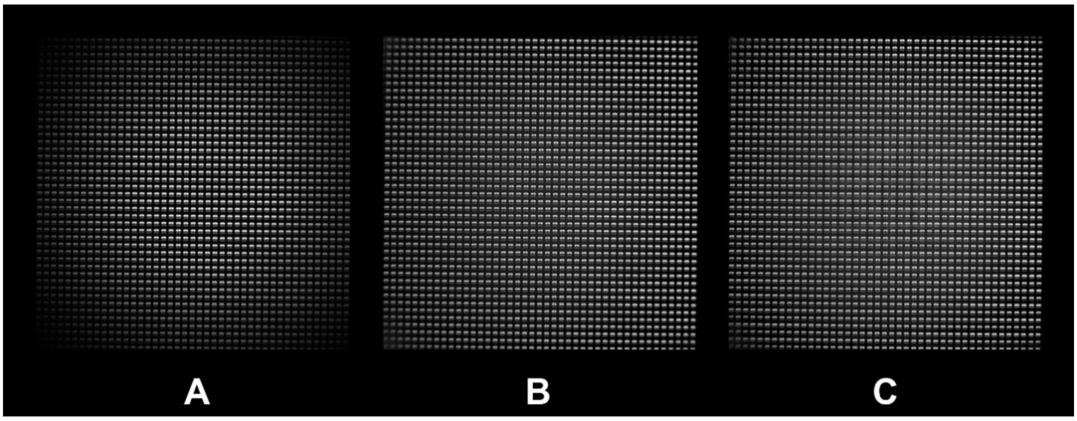


Figure 12. Test of multiscale flat field correction. (A) raw image obtained from the LCD screen of a smart phone imaged with the 2X objective. (B) Flat field correction obtained using the Laplacian pyramid. Flat field correction obtained from the Symlet wavelet method.

The method was subsequently used on experimental data (Figure 13). Different methods were tested. The seam observed at overlapping regions was most visible when blending using only the arithmetic average (Figure 13 A). Arithmetic averaging of the Laplacian pyramid method resulted in a more gradual transition in pixel intensity but long range variations could still be observed (Figure 13 B). Application of the blend exposure function further reduced the magnitude of the seam (Figure 13 C). The combination of flat field correction and Laplacian pyramid blending contributed to removing of discontinuities at the seam. Even the simplest blending method using arithmetic average improved on previous results (Figure 13 D). Performing image blending using the blendexposure removed all discontinuities (Figure 13 E).

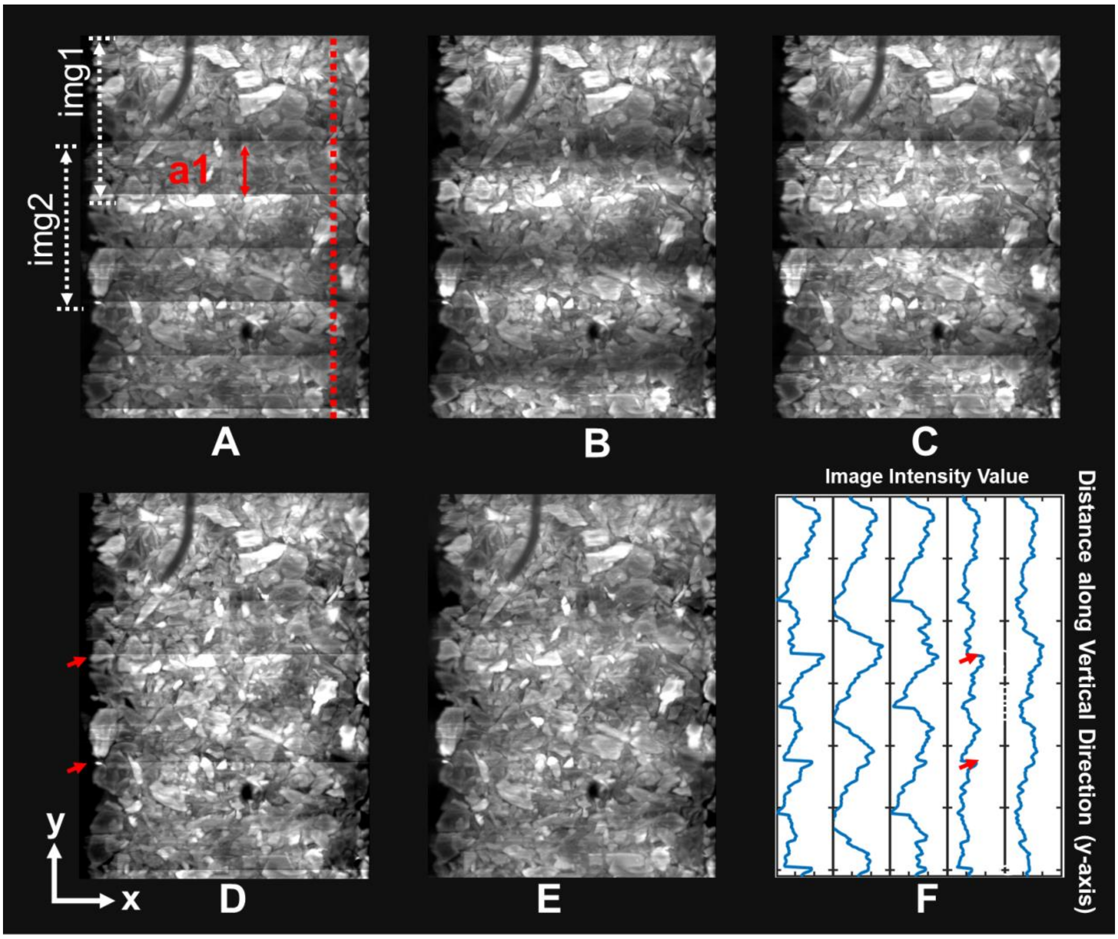


Figure 13. Testing of algorithms for stitching image data. (A) Stitching images using arithmetic average blending results in discontinuity where images overlap. a1 is the region of overlap of image 1 and image 2. The red line indicate the location where the image intensity profile is shown in (F). (B) Images stiched using Laplacian pyramid with arithmetic average. (C) Images stiched using Laplacian pyramid with blend exposure function. (D) Images stiched using Laplacian pyramid combining flat field correction and arithmetic average. (E) Images stiched using Laplacian pyramid combining flat field correction and blendexposure function. (F) Image intensity profiles along the vertical axis for image A to E (respectively from left to right).

**4 - Segmentation of image data**

Image segmentation was used to quantify the distribution and spatial arrangement of roots, soil particles and bacteria. Following segmentation, two important metrics were calculated, the 3D distance map and the map of the pore size. The following section describes the image segmentation pipeline developed to perform such measurements (Figure 14).

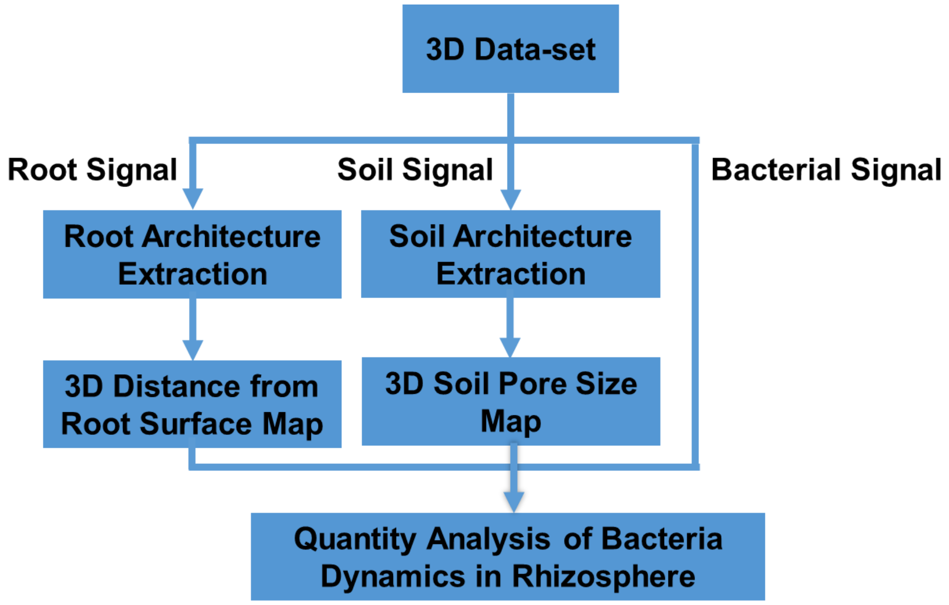


Figure 14 Image segmentation pipeline for the calculation of metrics characterising the location of bacteria with regards to the root and within the pore space.

The root domain was segmented from the light scattering signal using a region growing algorithm implemented in MeVisLab [8] following 6 steps.

1. A 3D image data is imported, filtered with a median filter, and visualised in a volume viewer.
2. In the transverse cross-section view (xz plane), one seed is placed at the base of the root system. When roots exhibit variations in contrast, a few additional seeds are placed along the root.
3. The region growing algorithm is run using a 3D 6-connect neighbourhood and with the lower threshold set above the mode value of the image histogram and with no upper threshold.
4. Steps 1 to 3 are repeated for the different time points using the same seed position. Adjustment of the seed positions and thresholding value is occasionally needed due to root movements and changes in scattering intensity of the root.
5. The resulting data is processed by the Euclidean Distance Transform function with 16-bit precision and the resulting volume data exported from MeVisLab for further processing.
6. Finally, the 3D distance map is incorporated into pipeline for quantification of bacterial cell density distribution.

The architecture of the soil particle was obtained using thresholding and morphological operators (Figure 15). The threshold $Thres_s$ is chosen marginally smaller than the mode value of the image (5% to 10% of the mode value of the histogram is usually adequate). This segmentation is not sufficient to extract out the core of soil particle (Figure 15 A), because the stain (sulforrhodamine-B) mainly attaches to the surface of the soil particles. Therefore, the segmentation approach also combined the GFP signal from bacterial fluorescence, using a different threshold $Thres_b$. The value of $Thres_b$ is choosen just above the noise level in the image (1 to 5% below the mode value of histogram is adequate). Median filtering with the window width set to the size of the soil pore is applied after thresholding (Figure 15 B). The following processes are used to segment the soil pore volume.

1. Determine $Thres_s$ from histogram of the red fluorescence image data ($I_s$)
2. Determine $Thres_b$ from histogram of the green fluorescence image data ($I_b$)
3. Apply media filter to $I_b$ to remove speckle noise.
4. Apply segmentation of soil particles using the following equation,

$$segmented\ soil\ (x,y,z) = \begin{cases} 1, & 0 < I_b(x,y,z) \leq Thres_b \\ 1, & Thres_s < I_s(x,y,z) \\ 0, & otherwise \end{cases} \quad . \qquad (12)$$

5. Apply morphological operators, i.e. close and erosion followed by fill operators (Figure 15 B b3).
6. Generate the structure of the soil pore using the inverse of the image generated in 5 and apply a mask to remove the padding domain.
7. Apply the local thickness metric [9,10] to determine the pore size distribution in the image. The algorithm was implemented as an ImageJ macro run in batch mode.

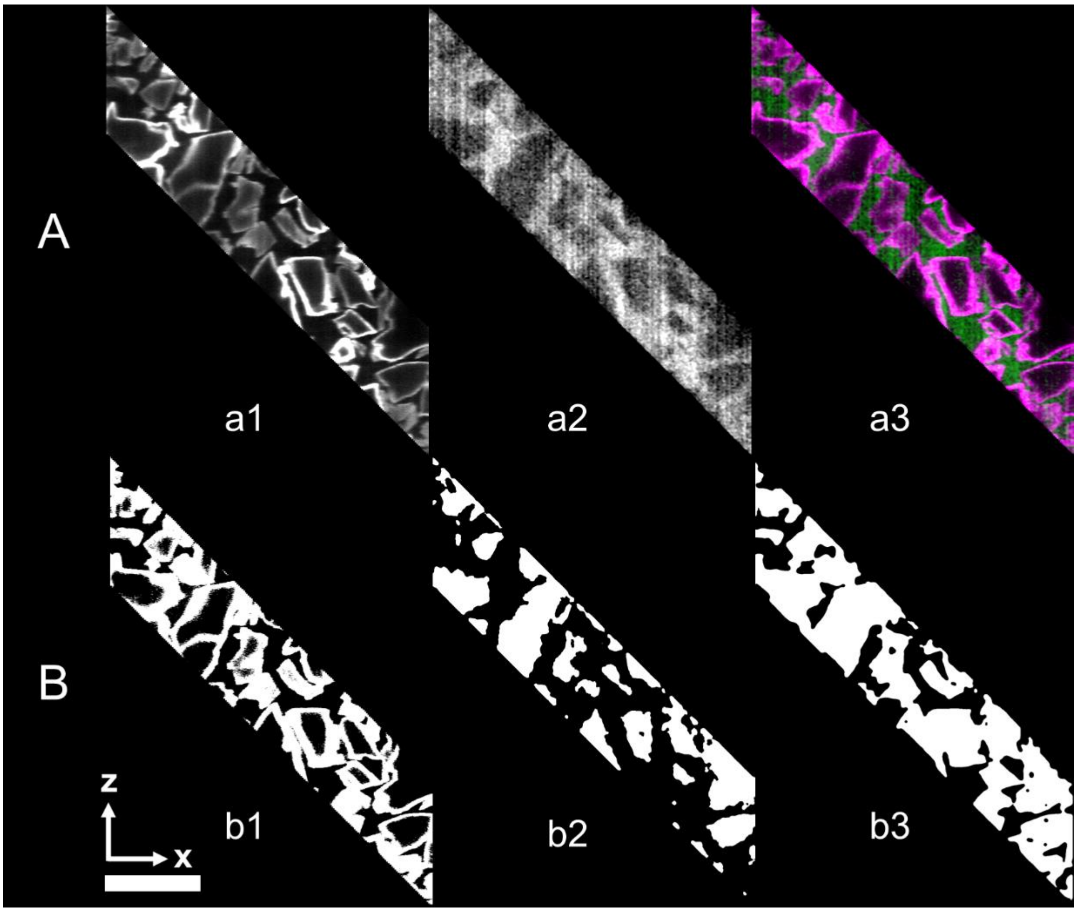

Figure 15. Segmentation of soil particles. (A) Cross section of fluorescent signals from bacteria and soil particles. (a1) fluorescent signal from soil particle, (a2) fluorescent signal from bacteria and (a3) overlay of bacteria and soil particle signals. (B) Segmentation of the pore space. (b1) thresholding of soil particle signal is followed by (b2) thresholding of the signal from the bacterial fluorescence signal. (b3) The segmentation of the interior of the soil particles is obtained by combining b1 and b2 (equation 12). Scare bar is 2 mm.

**5 - Quantification of bacterial density**

This section describes the calibration of the fluorescence intensity signal for the prediction of cell density from image data. Dense GFP-tagged *B. subtilis* cell suspensions were prepared in Percoll® and measured by OD600. Suspensions at ODs of 1.2 $10^{-3}$, 2.5 $10^{-3}$, 5.0 $10^{-3}$, 7.5 $10^{-3}$, 1.0 $10^{-2}$, 3.0 $10^{-2}$, 1.0 $10^{-1}$, 2.5 $10^{-1}$, 3.0 $10^{-1}$, 5.0 $10^{-1}$, 1.0 $10^{0}$, 2.0 $10^{0}$ and 3.0 $10^{0}$ were obtained by multiple dilutions. One ml of each bacterial suspension was transferred into a microscopy chambers and stained soil particles were added to adjust the focus of the microscope. The field of view of the microscope was then adjusted 2 mm above the soil particles. A full scan was obtained with 488 nm excitation illumination and GFP band pass emission filter, in steps of 50 µm and at two vertical positions 4 mm apart. The pixel intensity recorded in the images was then correlated to OD values, and OD values were subsequently correlated to bacterial counts (CFU). Estimation of bacterial cell density from image data were based on the resulting correlation (Figure 16 A), which showed a positive linear relation. The correlation could then be used to quantify bacterial cell density in live image datasets (Figure 16 B).

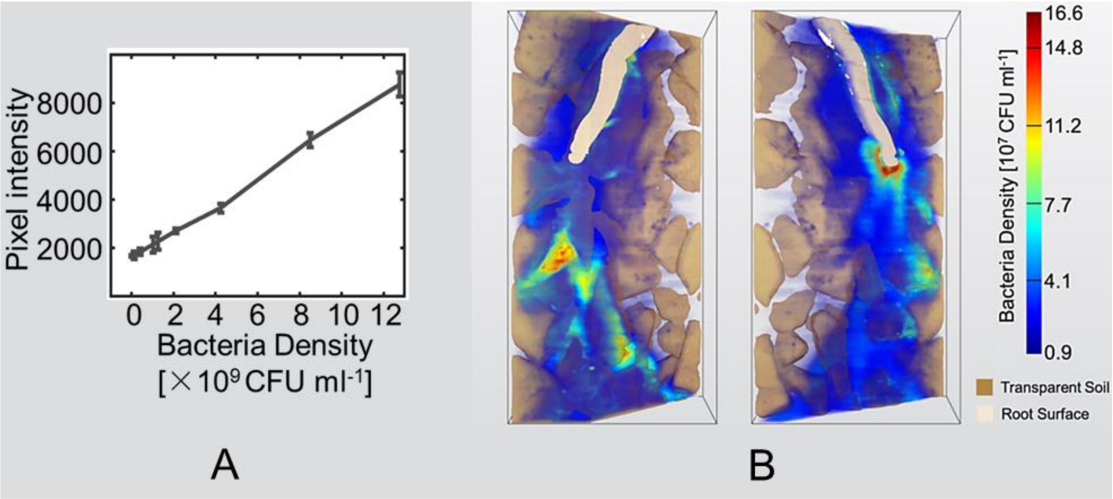


Figure 16. Quantification of bacterial cell density. (A) Relationship of *B. subtilis* cell density (CFU $ml^{-1}$) and pixel intensity (±SE). (B) Volume redering of bacterial cell density distribution in rhizosphere based on the the calibration of pixel intensity.

### 6 – Literature cited

8. MeVis Medical Solutions AG, F. M. MeVisLab. (2018).

9. Hildebrand, T. & Rüegsegger, P. A new method for the model-independent assessment of thickness in three-dimensional images. *Journal of Microscopy*

**185**, 67–75 (1997).

10. Dougherty, R. & Kunzelmann, K.-H. Computing Local Thickness of 3D Structures with ImageJ. *Microscopy and Microanalysis* **13**, (2007).
